# GOFlowLLM - Curating miRNA literature with Large Language Models and flowcharts

**DOI:** 10.1101/2025.10.07.680945

**Authors:** Andrew F Green, Nancy Ontiveros, Isaac Jandalala, Simona Panni, Valerie Wood, Giulia Antonazzo, Helen Attrill, Alex Bateman, Blake Sweeney

## Abstract

The exponential growth of non-coding RNA research—with over 230,000 papers published since 2000—has created an urgent knowledge management crisis in molecular biology. Despite their crucial regulatory roles, microRNAs (miRNAs) face a significant curation bottleneck, with only 1,400 articles manually curated to the Gene Ontology (GO) knowledgebase over a decade. We present GOFlowLLM, an automated curation pipeline powered by reasoning-enabled Large Language Models (LLMs) that follows established GO curation flowcharts to extract and structure miRNA-mediated gene silencing data at scale. When evaluated on existing curation, GOFlowLLM selects the correct GO term in 90% of cases. Curators also agree with 95% of the system’s reasoning steps and 90% of the evidence selected. Applied to 6,996 previously uncurated articles, our system identified 2,538 new candidate GO annotations on 1,785 articles in just 58 hours—potentially doubling the available miRNA GO curation. Manual review of a subset of these annotations shows that curators agreed with the selected term in 87% of cases, the model’s reasoning in 92% of cases, and the extracted evidence in 93%. GOFlowLLM demonstrates how LLMs can significantly accelerate biocuration while maintaining high-quality standards by following expert-designed reasoning frameworks. The integration of reasoning traces in our system provides transparent justification for annotations that can be reviewed by human curators, addressing one of the key challenges in adopting AI for scientific curation, potentially transforming how we manage the growing corpus of scientific knowledge in molecular biology. GoFlowLLM is available on github: https://github.com/RNAcentral/GO_Flow_LLM.

## Main

The exponential growth of non-coding RNA (ncRNA) research - over 230,000 papers since 2000, more than half of those since 2018 - has created a knowledge management crisis in molecular biology. ncRNAs regulate gene expression and participate in a wide range of biological processes, yet literature curation lags behind the pace of discovery. Most literature curation relies on only 100-200 biocurators^1^ who manually process scientific literature into structured knowledgebases like the Gene Ontology (GO). For microRNAs (miRNAs), limited resources have led to only 1,400 articles being added to the GO knowledgebase—a small fraction of the available literature, widening the gap between discoveries and structured knowledge. Few databases collect microRNA-target interactions, limiting access to regulatory networks. The IntAct team has applied controlled vocabulary developed for protein interactions to annotate ncRNAs, but the coverage remains limited^2^. MirTarBase^3^ is the most comprehensive microRNA database with thousands of annotations; however, most derive from high-throughput experiments. Results can be filtered by experimental evidence, such as luciferase assays, but even filtered subsets show low accuracy^4^. Similarly, RNAInter^5^ collects data through literature mining, with filtering options. None of these resources focus on microRNA action mechanisms, or directly link microRNAs to biological processes involving target genes, as GO annotations do. This curation bottleneck impedes scientific progress, as researchers struggle to identify the current knowledge and design boundary-pushing experiments. Our work addresses this challenge by developing an automated curation pipeline powered by reasoning-enabled Large Language Models (LLMs) that can accurately interpret and structure miRNA-protein interaction data from scientific literature at scale.

Biological knowledgebases are key infrastructure in modern biological research, providing integrated, reusable repositories of scientific knowledge. The Gene Ontology (GO) represents one of the most ambitious efforts, aiming to curate comprehensive gene function knowledge across the tree of life. Despite this broad mission, GO annotations remain heavily skewed toward well-studied organisms, particularly human^6^. This concentration is more acute for ncRNA, where annotation is carried out by a tiny fraction of biocurators, focusing almost entirely on *D. melanogaster* and project-based annotations to human, mouse and rat.

Previous attempts to accelerate GO literature curation have included text mining, natural language processing, and rule-based extraction tools^7,8^. While showing promise, these approaches struggle with scientific literature complexity, requiring integration of information across sentences, interpretation of specialized terminology, and nuanced judgments about experimental evidence in figures and graphs.

The GO consortium developed flowchart-based guidance for miRNA-mRNA functional interactions in post-transcriptional gene silencing to standardize manual curation^9^. This data type suits flowchart-based approaches since experiments and their interpretation are somewhat standardised - this extends to some other ncRNA types (lncRNA and siRNA, for example), potentially enabling automated curation for these important ncRNA types.

Our approach uses reasoning-trained LLMs to follow GO consortium flowcharts, providing an effective way to accurately annotate thousands of articles. This transforms annotation from human judgments into structured reasoning paths followed algorithmically while maintaining quality. The LLM provides specific article excerpts supporting decisions at each flowchart node, a novel feature in standard curation workflows outside of benchmarking exercises. We call this method GoFlowLLM.

GOFlowLLM follows current curator guidance (figure 1 flowchart) to curate three miRNA-mediated gene silencing terms: a general term, GO:0035195 (*miRNA-mediated post-transcriptional gene silencing*); and two more specific terms, GO:0035278 (*miRNA-mediated gene silencing by inhibition of translation*) and GO:0035279 (*miRNA-mediated gene silencing by mRNA destabilization*). Although tested only with the microRNA annotation flowchart and existing annotations, GoFlowLLM can be applied to any curation process modeled as a flowchart.

**Figure 1:**
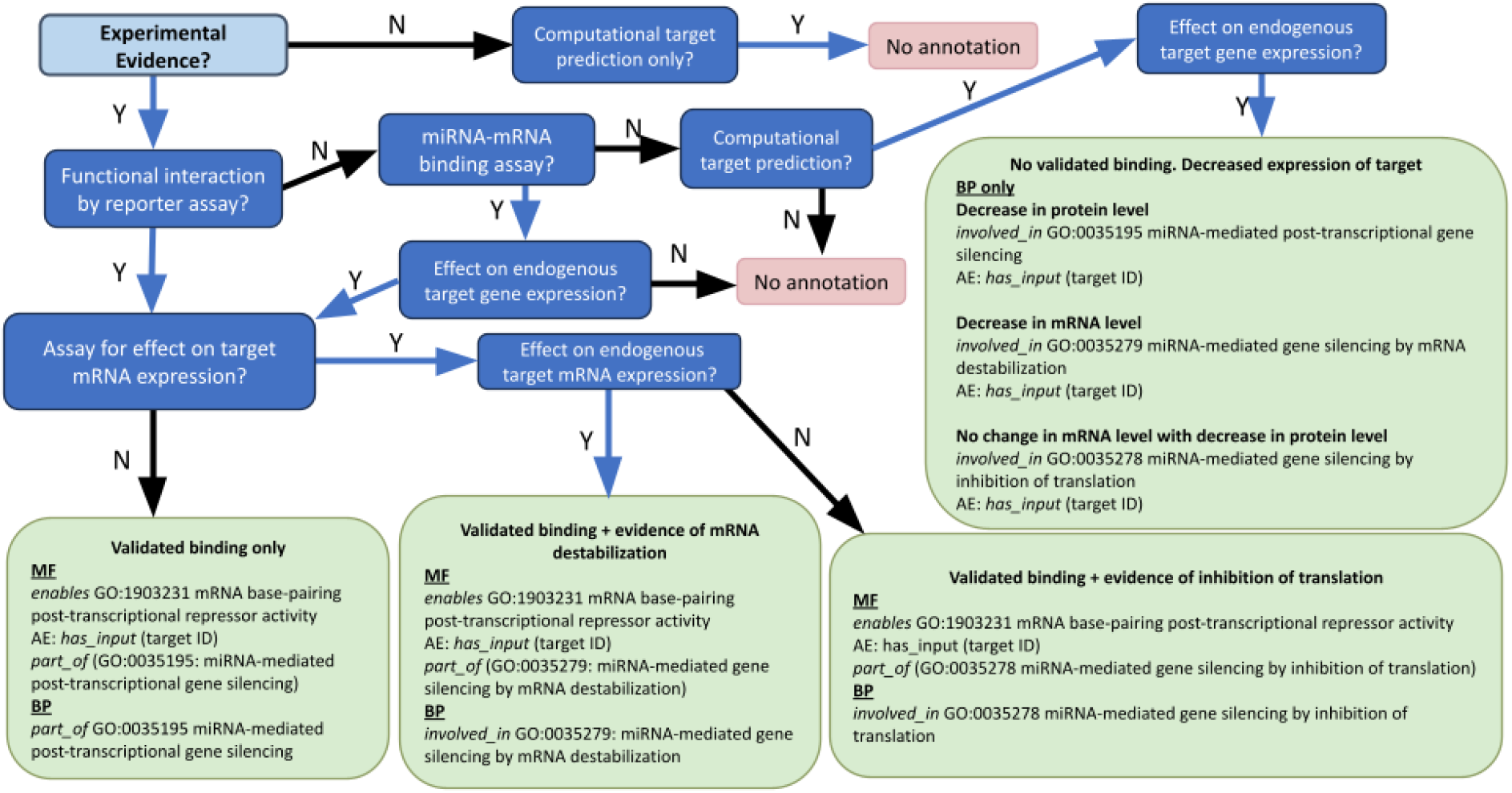
The microRNA flowchart used to curate microRNAs to the relevant GO terms of GO:0035195 (miRNA-mediated post-transcriptional gene silencing), GO:0035278 (miRNA-mediated gene silencing by inhibition of translation) and GO:0035279 (miRNA-mediated gene silencing by mRNA destabilization). This flowchart is an updated version of the original guidance published by the GO consortium in Huntley et al.^9,10^

GoFLowLLM was developed on sets of previously curated articles. On articles curated with recent guidelines (since 2022) GOFlowLLM agreed with manual curation in 4/5 cases. On articles curated with historical guidelines, GOFlowLLM reproduces the manually curated terms in 3/10 (30%) of cases, but upon re-curation is correct in 9/10 (90%) of cases, highlighting the potential for this tool to improve historical curation. In the 20 papers from the development, historical and recent annotations, curators agreed with 19/20 (95%) of the reasoning, 18/20 (90%) of evidence and 18/20 (90%) of both the evidence and final annotations. Targets were successfully extracted in 15/20 (75%) of cases.

We evaluated GOFlowLLM on 6,996 articles mentioning miRNA identifiers, deriving 2,538 new candidate GO annotations from 1,785 papers. These annotations were produced in 58 hours (2.4 days), equivalent in volume to all previous miRNA curation, covering numerous species without prior annotation. Curator evaluation on a subset of papers (60) show 86.7% correct annotations, with target and evidence extraction correct in 95% and 93.3% of cases, respectively. Combined with Argilla, GOFlowLLM creates a powerful curator assistant system automating candidate annotation production and simplifying human review.

## Results

We developed GOFlowLLM using curator guidlines and a small sample of 5 open access papers. A small number of papers were used for prompt development because curated papers are limited, and we needed sufficient papers for independent evaluation. During prompt development, GOFlowLLM produced correct annotations judged by human evaluation, but these did not always match existing annotations (Table 1). Investigation revealed that in late 2021, a new GO term, *miRNA-mediated gene silencing by mRNA destabilization* (GO:0035279), was added as a curation option. We hypothesised that this change to correspond to a change in the GO terms used in 2021, so we analyzed GoFlowLLM performance on two further publication sets: those with annotations made before 2021 (historical set) and annotations made after 2021 (recent set). We asked two questions; does GOFlowLLM reproduce existing GO annotations (Reproduction Accuracy)?; does it accurately produce correct annotations judged by expert curators using current guidance (Correctness)?

**Table 1:** Performance of GOFlowLLM on tested data sets. The low reproduction accuracy in the historical curation set is expected due to a change in the GO terms used in the curation flowchart, as described in the text.

| Dataset | Total Number of Papers | Manually Evaluated Papers | Reproduction Accuracy (%) | Correctness (%) |
| --- | --- | --- | --- | --- |
| Development (2015-2025) | 5 | 5 | 3/5 (60%) | 5/5 (100%) |
| Recent curation (2022-2025) | 42 | 5 | 4/5 (80%) | 4/5 (80%) |
| Historical curation (2015-2021) | 110 | 10 | 3/10 (30%) | 9/10 (90%) |
| Novel | 1,785 | 60 | NA | 52/60 (86.7%) |

**Table 2:** The number of annotations made to each of the three GO terms under consideration, and the observed failure modes of curation. Most failures occur because GOFlowLLM misattributes the results of an assay, usually qRT-PCR.

| GO Term | Total Annotations | Correct Annotations | Failure modes |
| --- | --- | --- | --- |
| GO:0035195 | 8 | 8 | - |
| GO:0035278 | 10 | 5 | Misattribution of qRT-PCR results, confusion over large number of assays, missing evidence of mRNA level change, out-of-scope biology |
| GO:0035279 | 42 | 39 | Misattribution of qRT-PCR experiment results, bad inference from multiple experiments |

After determining the performance of GoFlowLLM on a set of previously curated literature, and completing the first round of curator evaluation, we then applied GOFlowLLM to novel papers and evaluated correctness. When GoFlowLLM produced incorrect answers, we determined the type of error. This production run generated 2,538 new annotations on 1,785 previously uncurated papers.

We define accuracy as an exact match on the final GO term annotated by GOFlowLLM. Due to GO’s hierarchical nature, annotation to the less specific term (GO:0035195) remains technically ‘accurate’ even when more specific terms (GO:0035278 or GO:0035279) are possible. We use “accuracy” to refer to exact match accuracy for simplicity, as GOFlowLLM never selects the less specific term when a more specific one would be appropriate. We further distinguish between reproduction accuracy, and correctness; where reproduction is the ability of the system to reproduce the existing GO annotation, and correctness is the ability of the system to produce a correct GO annotation when judged by curators.

### LLM curation correctly reproduces human curation

We ran GOFlowLLM on all 42 articles annotated from 2022 onwards and compared with existing GO annotations. The existing GO term was reproduced in 34/42 papers (81%). Manual evaluation of 5 sampled articles found 4/5 (80%) reproduced existing annotations and were correctly annotated. The one incorrect annotation involved misattribution of qRT-PCR results leading to overly specific annotation; the curator recognized this as a confusing case.

### LLM curation can update outdated annotations

We ran GOFlowLLM on 110 historical papers annotated before 2022, where the existing GO term was reproduced in only 25/110 papers. We identified the reason for this poor reproduction accuracy as a change in the curation guidelines in late 2021, where a more specific term was introduced: historical annotations are still correct, but can be made more precise. Manual evaluation of 10 sampled papers showed poor reproduction accuracy (3/10) but excellent correctness (9/10). The one mistake involved the same failure mode: misattribution of qRT-PCR results causing overly specific annotation. This performance shows GOFlowLLM can review historical annotations to make them compliant with current guidelines, and deepen the annotation for existing literature. It confirms our hypothesis that poor reproduction accuracy reflects changing guidelines, not tool failure.

GO miRNA curation evolution can be traced through the GO annotation guideline wiki pages ^11^ revision history, where more specific ontology terms have been introduced with detailed evidence requirements. Since re-curation typically addresses incorrect annotations rather than less specific ones, annotations that could be improved by assigning a more specific new GO term are often missed in data retrofits. This highlights GoFlowLLM’s potential for rapidly improving existing annotations when GO standards change.

### GoFlowLLM identifies 2,538 novel annotations from 1,785 unseen papers

We next used RNAcentral’s LitScan tool to identify 6,996 open-access peer-reviewed articles mentioning one miRNA using miRBase and MirGeneDB identifiers^12,13^. These articles were analysed by GoFlowLLM providing GO annotations for 1,785 papers. Un-annotated papers contained experiments that were not suitable for the GO annotation in the miRNA curation flowchart (e.g. differential expression analyses, in-silico target prediction, pathway analysis, etc), these articles were filtered from subsequent stages. Many of the 1,785 papers contained multiple miRNA targets providing miRNA-mRNA interaction annotations for 1,024 miRNAs to 2,538 target mRNAs.

To assess annotation correctness, we randomly sampled 60 papers for manual review. Of these 52/60 (86.7%) were correct, closely matching the accuracy of re-annotated historical curation (90%).

We observed the same failure modes as in the re-annotated sets: mostly qRT-PCR misattribution leading to overly specific terms. However, the broader literature pool revealed other failure modes, including the inclusion of out-of-scope biology, missing evidence, and confusion caused by papers with numerous assays. Some cases mentioned the mandatory qRT-PCR experiment, but only reported it fully in the supplementary materials, which GOFlowLLM cannot access. These suggest improvement areas for both our prompts and the curation flowchart itself, making evidence requirements and in-scope experiment types more explicit.

GOFlowLLM has annotated articles covering interactions across many species, contrasting human curation’s focus on model organisms. Despite this, GOFlowLLM more than doubles available curated literature in major model organisms and provides coverage for otherwise uncurated species. Figure 2 shows the taxonomic distribution comparison of existing GO annotations for the three miRNA binding terms.

**Figure 2:**
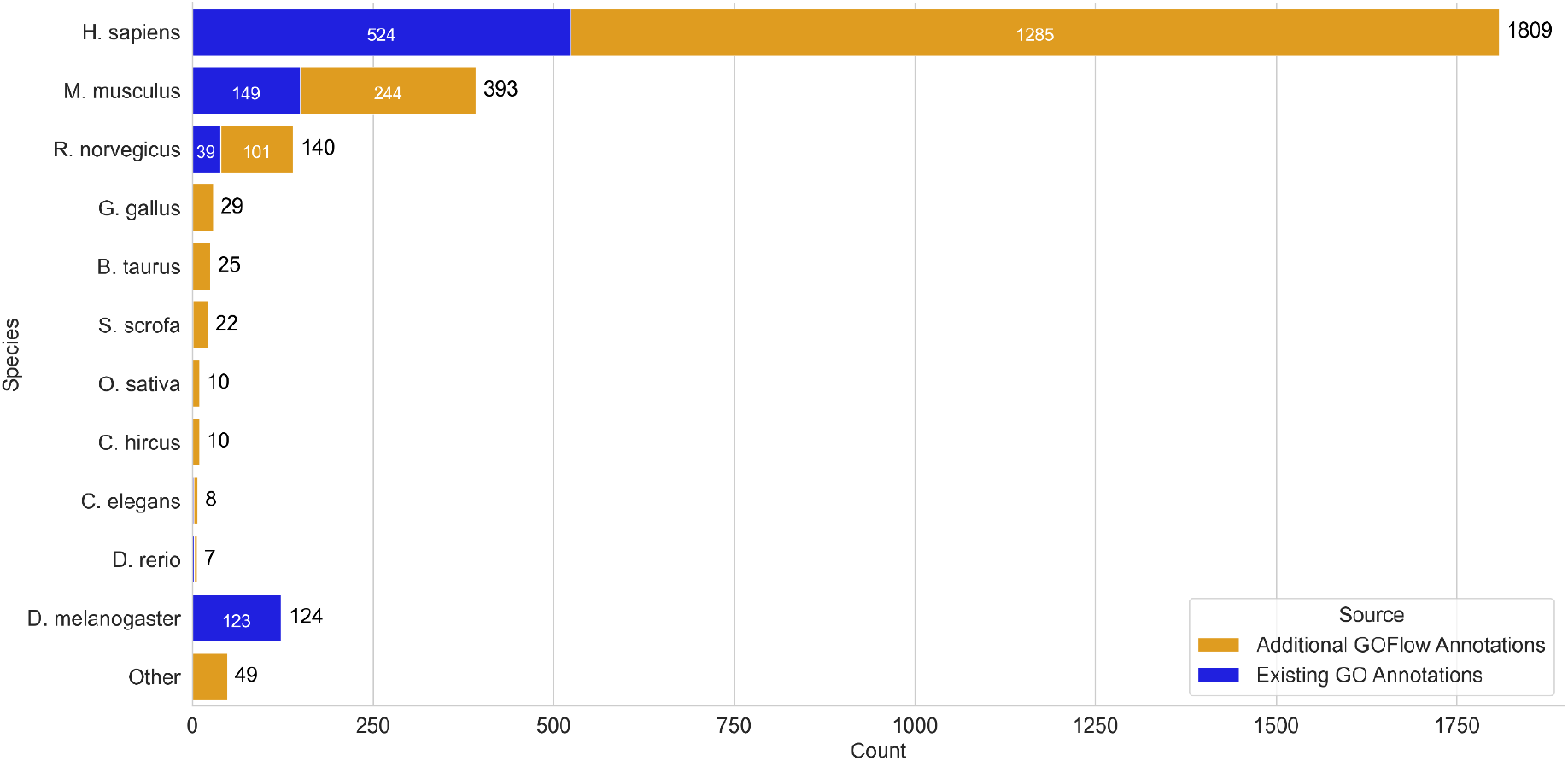
Bar chart comparing the taxonomic distribution of GoFlow LLM annotations to existing GO annotations for the three terms annotated. It is important to note that the papers with existing GO annotation include closed source literature and open access, while GoFlowLLM’s annotations are only on open access literature, and restricted to papers mentioning only one miRNA with a species specific identifier. The limited number of additional D. melanogaster identified is due to recent efforts at FlyBase to curate all miRNA gene silencing experiments.

Of the annotated articles, 118 (6%) discuss organisms lacking existing GO annotations for miRNA mediated gene silencing terms, increasing GO species coverage. Notably, while the existing GO annotation distribution includes both open and closed-access articles discussing any number of miRNAs, GOFlowLLM is restricted to open-access articles with single miRNAs. The limited number of additional *D. melanogaster* identified (figure 2) is due to recent efforts at FlyBase to curate all miRNA gene silencing experiments^14^.

GOFlowLLM averages 109 seconds per article across 6,996 articles, equating to approximately 790 papers per day. Using multiple processes on 4xA100 GPUs, we processed the entire article set in 58 hours (2.4 days).

### GoFlowLLM annotations use appropriate good reasoning and evidence

Curator assessment used Argilla^15^ to display full LLM traces (reasoning, decision and evidence from the context window). Across all 20 papers from the sets labelled development, historical, and recent, curators agreed with 19/20 (95%) reasoning traces, 18/20 (90%) evidence extractions, and 18/20 (90%) of final annotations. Targets were successfully extracted in 15/20 (75%) cases.

The tool selects accurate evidence, with curators judging 90% of annotations well-supported. Most failures occurred because constrained generation substring selection could not access the article section containing the most relevant information - a limitation of the guidance toolkit, requiring section-by-section processing to avoid memory issues. Evidence selected for novel set annotations is adequate in 56/60 (93.3%) of cases. Most failures involve extracting somewhat relevant but suboptimal evidence, as assessed by the curators. In some cases, the most appropriate evidence does not exist in the expected paper section from which we extract substrings - a limitation of the toolkit ensuring evidence is a true paper substring.

Target extraction performed well, achieving complete target recovery in 15/20 articles. The most common failure (4/5 of cases) involved selecting too many mRNA names from the EuropePMC list; additional genes were mentioned in the paper, but not validated by a luciferase assay as miRNA targets. In one case, the LLM failed because the correct target was absent from the list provided by EuropePMC. In the 15 correct target identifications, GoFlowLLM extracted all validated targets mentioned, ranging from one to four targets per article.

GOFlowLLM’s curation report is a valuable artefact representing previously nonexistent curation records. While curators may note the specific supporting sentences, they would not produce detailed reports evaluating evidence and appropriate GO term selection.

## Discussion

### Comparison to existing tools

Integrating large language models into the biocuration workflow offers a promising solution to address the growing challenge of literature curation in the rapidly expanding field of ncRNA research. Our study represents the first demonstration of an LLM-driven curation tool specifically designed for ncRNA GO annotation, achieving high accuracy in reasoning and annotation assignment through expert curator evaluation.

Several LLM-based biocuration approaches are already being developed. SPIRES^16^ uses GPT-4 to extract structured information for gene-chemical-disease associations, while CurateGPT^17^ focuses on ontology curation and creation. More closely aligned with our approach are AutoMAxO^18^, which annotates rare disease literature using the Medical Action Ontology, and FuncFetch^19^, which extracts structured information about enzymes from literature. However, unlike these tools that target different domains, our GoFlowLLM tool specifically addresses ncRNA functional annotation—an area that has received less attention in biocuration efforts.

No published work specifically addresses LLM-based curation for ncRNA function, highlighting the novelty of our approach. The integration of reasoning traces in our system is particularly notable, as these provide transparent justification for annotations that can be quickly reviewed by human curators, addressing a key challenge in adopting AI for scientific curation.

### Limitations

Despite the promising results, our current implementation has several limitations that present opportunities for future improvement. First, our flowchart-based approach is designed to work with only one RNA at a time, which limits throughput when processing papers that discuss multiple RNA-target interactions. We have also not addressed the curation of miRNA clusters, which require a different approach as the cluster of miRNAs acts in concert to exert a regulatory effect.

The GO terms annotated by GoFlowLLM relate to only three specific Biological Process (BP) terms and one Molecular Function term—a tiny subset of the overall GO BP domain, which has around 26,000 terms. However, by constraining ourselves to this small subset, we ensure high accuracy annotation. GoFlowLLM serves as a curator assistant, identifying candidate miRNA-mRNA pairs and likely mechanisms, allowing curators to focus on less well-defined biological aspects that would be difficult to curate reliably with an LLM.

Relatedly, GOFlowLLM’s core design is around the use of a curation flowchart. This works very well for some aspects of GO curation where the required evidence is quite narrowly defined and a clear decision process can be drawn from the contents of a paper to the eventual GO term annotated. However, this is not representative of all GO curation. It is unlikely to be immediately applicable more broadly without significant effort, and will struggle to capture the curation process for novel biology.

While the quality of evidence extracted by GOFlowLLM is high (93.3%), there is room for improvement. We rely on the guidance library for constrained decoding to extract evidence guaranteed to be contained in the paper, but it cannot yet reliably select substrings from the entire paper. Additionally, the LLM sometimes overlooks evidence that would support more detailed annotation, partly due to loading only predetermined sections of papers.

Finally, we depend on the EuropePMC annotations API for gene symbols (i.e. their common names, like SMAD4), which occasionally provides incomplete lists leading to incorrect target identification. The LLM could identify correct symbols but was prevented from reporting them when they weren’t in the EuropePMC list.

### Broader impact & future directions

Our work aligns with the broader trend of leveraging AI to address the curation bottleneck in bioinformatics. By automating the initial stages of literature review and annotation, these tools allow human curators to focus their expertise on validation and integration of findings, potentially increasing both the throughput and quality of biocuration efforts. As LLM capabilities continue to advance, we anticipate that the accuracy and scope of automated curation will expand, gradually reducing the manual effort required for maintaining comprehensive and up-to-date biological databases, while expert curators remain essential for validation and ensuring biological coherence.

GOFlowLLM’s ability to provide more specific annotations than older curation represents a significant strength of the approach. We observed that GOFlowLLM diverges from older curation in ways that reflect evolving curation guidelines, with expert curators agreeing with 90-100% of GOFlowLLM’s revised decisions. This highlights the utility of LLM-based systems for retrofitting existing annotations with more specific terms when supported by evidence, whether for GO or other ontologies.

In future iterations, we aim to give the tool more agency—allowing it to determine which entities to annotate with which terms, rather than following a predetermined flowchart. Integration with external knowledge bases could improve curation accuracy for entities like cell lines and disease states that are often described using non-standard terminology. We also see potential in adapting this decision tree-based approach to other non-coding RNAs, such as lncRNAs and siRNAs, and extending beyond GO annotations to cover expression patterns, subcellular localization, and biomolecular interactions. By broadening scope while maintaining high accuracy, such tools could significantly enhance ncRNA databases and facilitate new discoveries in RNA biology.

### Conclusion

In conclusion, our LLM-driven approach represents the first automated tool specifically designed for ncRNA GO annotation, achieving high accuracy in both reasoning and annotation assignment as validated by expert curators. By providing transparent reasoning traces and enabling more specific annotations than legacy curation, GoFlowLLM offers a practical solution for addressing the biocuration bottleneck in RNA biology. While challenges remain in expanding scope and autonomy, our results demonstrate that LLM-assisted curation can significantly accelerate the integration of scientific findings into structured knowledge resources while maintaining the quality standards required for biological databases.

## Methods

### Curation flowchart

The latest version of the miRNA curation flowchart was downloaded from the GO wiki^11^ (figure 1), converted to JSON for GOFlowLLM, and prompts were prepared for each node in an accompanying JSON file.

The miRNA curation guidelines focus on mechanisms by which miRNAs silence gene expression by binding target mRNAs; as such, the number of terms considered is small, primarily those that define the mechanism by which the miRNA silences gene expression. The least specific term GO:0035195 (miRNA-mediated post-transcriptional gene silencing) has two child terms separated by the mechanism of silencing: GO:0035278 (miRNA-mediated gene silencing by inhibition of translation) and GO:0035279 (miRNA-mediated gene silencing by mRNA destabilization). Most existing annotations fall into the cases along the bottom of the flowchart, and will also have the MF term GO:1903231 (mRNA base-pairing post-transcriptional repressor activity) and a specific BP term that describes mechanism of gene silencing. Figure 3 shows the local hierarchy of these terms.

**Figure 3:**
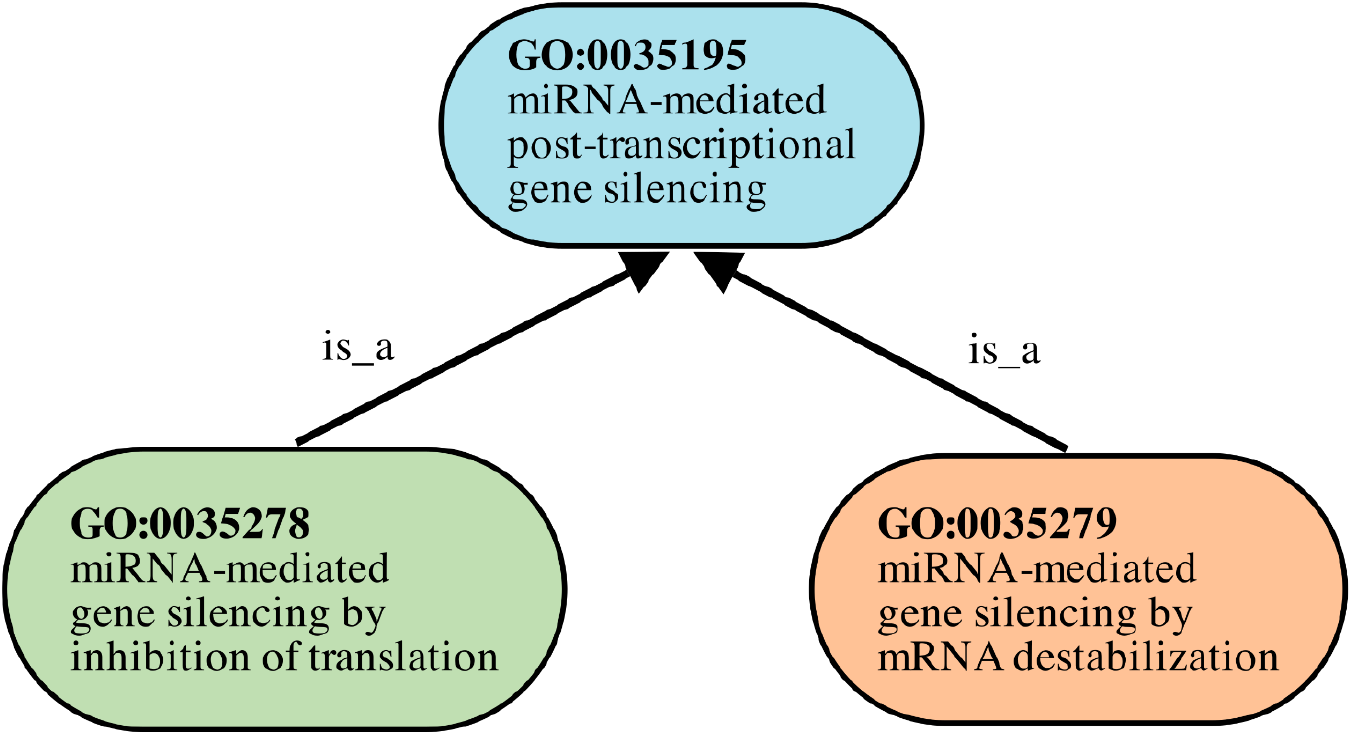
Local hierarchy of the three terms annotated by GOFlowLLM. The two more specific terms (GO:0035278 and GO:0035279) are child terms of GO:0035195, related in the ontology by the is_a relationship.

The term hierarchy is important when evaluating against existing curation, where less specific annotations may be ‘deepened’ to more specific terms by following current guidelines specifying the required experiments and experimental results.

### Dataset preparation

GO annotation files for ncRNA are retrieved from the GOA project at EBI as a GPAD file, and processed to select ncRNAs annotated by the UCL BHF project^20^. Of 448 unique publications, 157 are open access with full text available via the EuropePMC API. The annotations were made over approximately 10 years, (figure 4) and include GO terms ( GO:0035195, GO:0035278 or GO:0035279), interacting miRNA (RNAcentral URS_taxid), and target protein (UniProtKB accession ID). We converted the URS_taxid and UniProtKB accessions to human-readable identifiers using lookup tables linking URS_taxid to standard RNA ID, and UniProtKB accessions to protein symbol.

**Figure 4:**
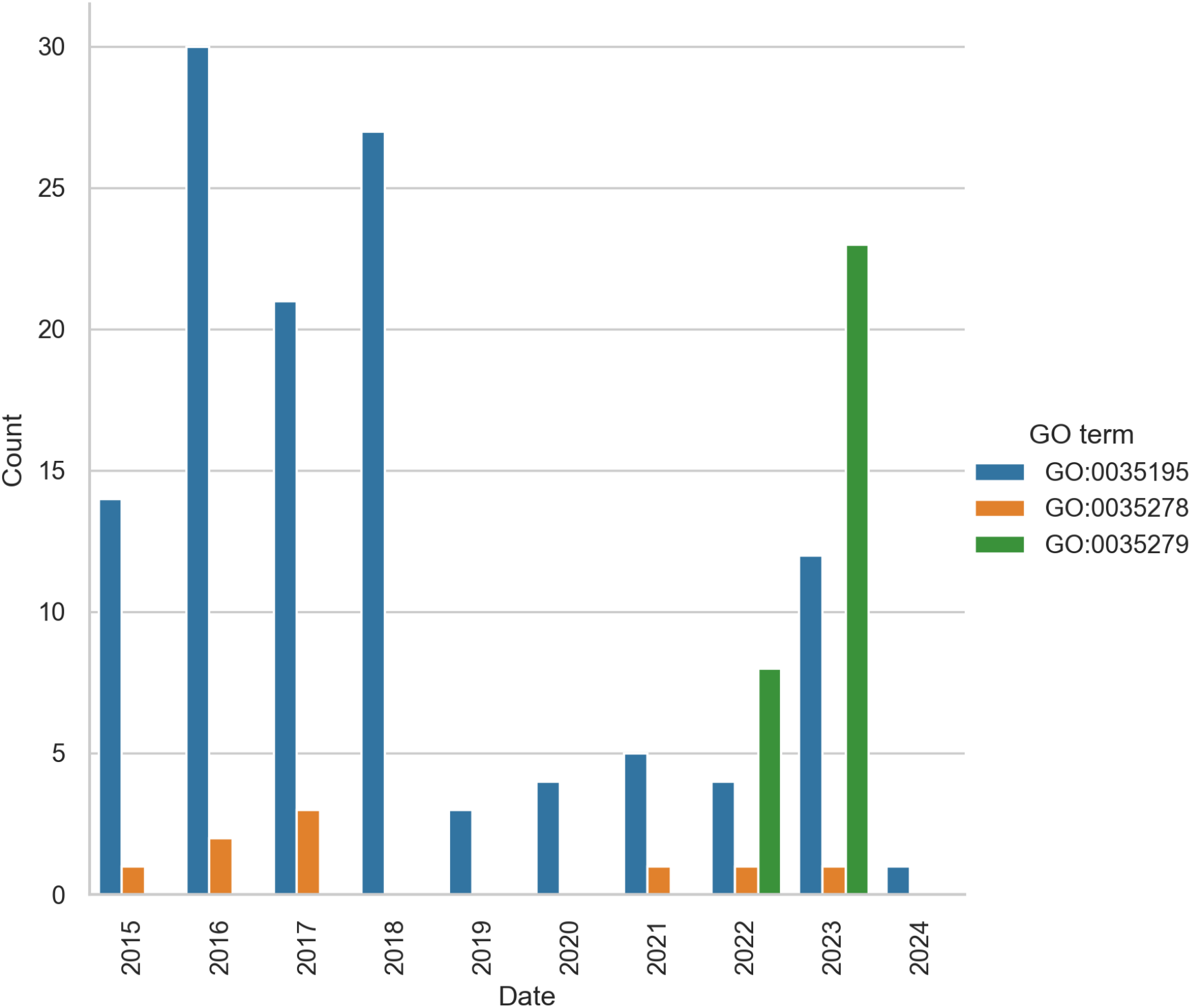
Annotations to each GO term since 2015, restricted to only open-access publications. The majority of annotations were made in 2016-2018, with GO:0035195 being the most common term annotated.

The annotations were produced using a variant of the flowchart from Huntley *et al*.^9^. Since curator decision records do not exist, multiple plausible flowchart paths often exist for a given annotation. We therefore assess our system using only the final annotation and record decisions at each flowchart node for review, rather than retrospectively inferring flowchart paths from annotations alone. Since all GOFlowLLM annotations require human curator review before GO inclusion, the combination of GOFlowLLM and the Argilla tool we use for human evaluation creates a powerful curation assistant.

We split 157 open access papers into three sets: 5 papers for prompt development with curator guidance; 110 papers annotated pre-2022 using older guidelines labelled as “historical”; and 42 papers annotated since 2022 labelled as “recent”. We ran GoFlowLLM on all papers, and sampled 10 historically curated papers and 5 recently curated papers for detailed manual review, assessing reproduction accuracy, annotation correctness, reasoning correctness and evidence usefulness.

To assess broader literature curation utility, we collected publications mentioning miRNA using RNAcentral’s LitScan tool. This tool searches the EuropePMC API for ncRNA identifiers and retrieves PMIDs and sentences to build a picture of the literature around a given ncRNA. The tool is described in more detail in Green *et al*.^21^. One key limitation is that LitScan, and consequently our tool, only analyses open access literature.

To minimise false positives, we restricted miRNA IDs to those from miRBase and mirgenedb^12,13^, considered authorities on miRNA naming with organism-specific identifiers. Detecting these identifiers strongly indicates that the miRNA studied matches the database and can accurately be linked to a URS taxid. We further required articles to mention only one miRNA and excluded retracted and review articles, which are unsuitable for GO curation. This provides a set of 6,996 articles for potential curation.

### LLM framework

We use the QwQ-32B model from Qwen^22^, which has 32 billion parameters and supports 128k token context length, making it ideal for this application. While QwQ is not the state-of-the-art for local reasoning models, it runs with fewer resources at higher precision and higher speed; an acceptable tradeoff when processing thousands of publications. For efficiency, we use a quantized version with 8-bit integer weights, improving performance with negligible quality degradation. We run the LLM on Nvidia A100 GPUs with 80GB VRAM restricting the context to 64k tokens to stay within single-card VRAM limits.

Since our LLM follows a flowchart, it must reliably answer questions in a predetermined format - single words ‘yes’ or ‘no’. Ordinarily, this requires strong prompting and post-processing, but constrained decoding offers a more efficient approach. In constrained decoding, we alter the available token space during decoding, sampling only tokens fitting our requirements. This technique forces the model to choose between ‘yes’ and ‘no’ without ambiguity, eliminating post-processing. We use the guidance framework^23^, and run the LLM using the widely used llama.cpp tool^24^. Guidance also controls generation parameters like temperature and reliably stops generation at tokens of our choosing, or after a specified token count. One downside is that constrained decoding precludes API model use, since token sampling occurs behind the API - this excludes state-of-the-art models like OpenAI’s o-series or Anthropic’s Claude models.

To enable fast annotation scaling to thousands of papers, GOFlowLLM works on article sections rather than whole texts. This provides the section name most likely to contain each step’s answer and loads associated text only if not already been loaded. This minimises context while exploiting the LLM’s excellent recall for previously seen sections. For substring extraction, the predicted relevant section is provided the LLM to select from, while guaranteeing substrings come directly from the paper, guidance limits maximum text size for substring sampling to limit memory required and maintain performance. Due to this text limit, evidence extraction may not always extract the most relevant passage.

When loading article sections, we use standard labels like ‘Methods’ and ‘Results’. However, relying solely on these standard names is insufficient due to the wide variability in section headings across articles. We use zero-shot classification to ask the LLM to select the most appropriate heading from the article’s actual headings, based on our standard headings. For example, when searching for the ‘Methods’ section, the model may identify ‘MATERIALS AND METHODS’. This allows seamless handling of different article styles.

A key aspect of GO annotation is correctly identifying the miRNA’s target mRNA. While we guarantee the extraction of a valid substring, substrings may not always end at correct boundaries (including additional text). We reuse the EuropePMC annotations API^25^, which lists all genes/proteins identified in open access publications. The API endpoint we use to provide the structure we parse to extract gene names is:

~~~
https://www.ebi.ac.uk/europepmc/annotations_api/annotationsByArticleIds?articleIds=PMC:{pmcid}&type=Gene_Proteins&provider=Europe PMC
~~~

EuropePMC documentation provides additional details of the API^26^. Constrained generation then selects the most appropriate gene symbol from the list.

QwQ is a ‘reasoning model’, designed to generate reasoning tokens (a reasoning trace) before answering any questions. This model type excels at complex problems and allows variable inference-time compute by generating additional reasoning tokens. We prompt QwQ to produce up to 1024 reasoning tokens before answering. While relatively small for a reasoning trace, experiments showed the model rarely needs more tokens, and recent work indicates this should be adequate for our domain^27^. We retain reasoning traces in curation records to help identify logic failures that can be fixed through prompting.

We sought to extract the specific text sections supporting curation decisions. Currently, curation does not require this, with the granularity of supporting evidence usually at the article level. Guidance leverages the LLM to select text substrings and, with accompanying prompts, generate supporting statements for that choice. This allows extracting article substrings justifying corresponding curation decisions.

### LLM instruction and prompt development

The flowchart nodes are written as yes/no questions and serve as the starting point. We add further guidance when a question is ambiguous or greater selectivity is required. For example, the node “Functional interaction by reporter assay?” has the following prompt:

> *Does the paper have a functional interaction between the miRNA and an mRNA, determined by a reporter assay? Acceptable assays include: Luciferase reporter assays where a luciferase reporter gene is fused to the 3’UTR of the target mRNA; and CRISPR/CAS9 deletion of the miRNA response element with subsequent measurement of protein levels*.

This prompt incorporates the decision tree node question and includes GO miRNA curation manual guidelines (https://wiki.geneontology.org/MicroRNA_GO_annotation_manual). We deliberately restricted acceptable assays to the most widely used, requiring minimal additional reasoning and evidence.

To develop the prompts, we sampled 5 articles from previously curated articles, designed each prompt to correspond to the nodes in the flowchart with curator guidance. Prompts were manually tuned to arrive at the correct final annotation as judged by the curators - not necessarily matching historical annotation.

The flowchart incorporates filtering elements (e.g. checking experimental results presence). However, processing numerous articles required two additional filtering nodes before the existing decision tree to reduce forwarded articles. These exclude articles not mentioning binding experiments, which should not be curated for these GO terms, and miRNA cluster articles, which require different, more complicated treatment.

### Human evaluation

Expert curators validated GOFlowLLM output, considering model’s reasoning and the full text of the paper. The model’s reasoning, evidence and decision were loaded into the Argilla tool^15^ hosted at RNAcentral. We evaluated 5 articles from the recent set, and 10 from the ‘historical’ set.

Curators were asked the following questions:

- Does the reasoning from the model address the questions well?
- Does the provided evidence support the decisions given for the questions?
- Is the final annotation correct?
- Are all targets identified correctly?

Curators could also comment on the output and identify over-annotated or missed targets.

We sampled 60 papers from the broader literature annotations, loading them with the full trace of prompts, reasoning, evidence and decisions and final annotations. We use the same question set as above to evaluate performance on these unseen articles.

